# Low-coverage whole-genome sequencing facilitates accurate and cost-effective haplotype reconstruction in complex mouse crosses

**DOI:** 10.1101/2025.05.19.654056

**Authors:** Samuel J. Widmayer, Lydia K. Wooldridge, Emily Swanzey, Mary Barter, Chrystal Snow, Michael Saul, Qingchang Meng, Beth Dumont, Laura Reinholdt, Daniel M. Gatti

**Affiliations:** The Jackson Laboratory for Genomic Medicine, 10 Discovery Drive, Farmington, CT, USA; The Jackson Laboratory, 600 Main St, Bar Harbor, ME, USA

**Keywords:** mouse, Diversity Outbred, low-coverage WGS, QTL mapping

## Abstract

The search for the underlying genetic contributions to complex traits and diseases relies on accurate genetic data from populations of interest. Outbred populations, like the Diversity Outbred (DO), are commonly genotyped using commercial SNP arrays, such as the Giga Mouse Universal Genotyping Array (GigaMUGA). However, array genotypes are expensive to collect, subject to significant ascertainment bias, and too sparse to capture the genetic structure of highly recombined mouse crosses. We investigated the efficacy of sequencing-based genotyping by comparing genotyping results between the GigaMUGA, double-digest restriction-site associated DNA sequencing (ddRADseq), and low-coverage whole-genome sequencing (lcWGS). We aligned reads at ∼1X coverage and imputed segregating SNPs from the eight DO founder strains onto 48 DO genomes and reconstructed their haplotypes using R/qtl2. Haplotype reconstructions derived from all three methods were highly concordant. However, lcWGS more faithfully recapitulated crossover counts and identified more small (< 1 Mb) haplotype blocks at as low as 0.1X coverage. Over 90% of local expression quantitative trait loci identified in a set of 183 DO-derived embryoid bodies using the GigaMUGA were recalled by lcWGS at coverages as low as 0.1X. We recommend that lcWGS be adopted as the primary method of genotyping complex crosses, and cell-based resources derived from them because they are as accurate as array-based reconstructions, robust to ultra-low sequencing depths, may more accurately model haplotypes of the mouse genome that are difficult to resolve with dense reference data, and cost-effective.

## INTRODUCTION

The field of systems genetics relies on the integration of diverse molecular data types to understand the genetic pathways and molecular networks that underlie complex disease. Investigations in laboratory model systems have been at the forefront of systems genetics discovery since the field’s inception due to their experimental tractability and ready availability of diverse tissue. Two complementary resources for systems genetics in the mouse are the Collaborative Cross (CC) strains (Churchill et al., 2004), (Collaborative Cross, 2012), (Saul et al., 2019) and the Diversity Outbred (DO) mouse population (Churchill et al., 2012), (Chesler et al., 2016). Both populations were founded through organized breeding programs initialized from eight genetically divergent inbred strains and capture significant genetic variation that more accurately reflects human population diversity. The mouse community has invested substantial time and resources in making these mouse diversity resources as powerful and flexible as possible. This has been achieved in part by optimizing the process of obtaining high-quality genotypes via the Illumina Infinium-based Mouse Universal Genotyping Array (MUGA) series of microarrays, which were originally tailored to identify haplotype blocks in the CC and, more often, the DO (Morgan et al., 2015).

The MUGA arrays were developed to standardize and streamline genotyping from complex mouse crosses and to distinguish among the three cardinal mouse subspecies. Thus, the performance of many array markers from dozens of mouse strains is readily available and routinely used to quality check genotypes obtained in mouse crosses. Consensus genotypes for the eight founders of the CC and DO aid in genetic analysis with standard analysis tools like R/qtl2 (Broman et al., 2019). In addition, because the CC strains are almost entirely inbred, consensus genotypes for each CC strain also exist for sites represented on the GigaMUGA (Shorter et al., 2019). Together, these considerations make the GigaMUGA an attractive option for genotyping mice from complex crosses.

Clear limitations remain for studies using the DO and other complex crosses derived from genetically diverse mouse strains. First, whole-genome sequencing of CC strains attempted to fill marker genotype gaps and revealed *de novo* mutations that have accumulated in certain CC lines (Srivastava et al., 2017), demonstrating both the utility of direct, sequence-based characterization of already well-characterized genetic reference populations, and the potential pitfalls of continuing to rely on array genotypes. Second, detecting QTL of modest effect with high power requires between 200-500 DO mice (Keele, 2023). DO mice are non-fungible, and genotyping a mouse cohort on the GigaMUGA often represents a significant cost for investigators. Crosses between CC strains (CC-RIX) and outcrosses of various mutant strains to DO animals (DOF1) may identify segregating modifiers of notable mutant phenotypes (Saul et al., 2019), (Hackett et al., 2022), (Kim et al., 2023). However, to achieve optimal mapping power, these complex cross designs may require significantly greater sample sizes than ordinary DO or CC mapping experiments and, therefore, require a substantial genotyping investment. Third, quantitative genetic analysis using the DO and CC typically models the additive effect of each of the eight founder alleles at each marker, which requires founder haplotype reconstruction (Svenson et al., 2012). With each outbreeding generation in the DO, crossovers accumulate. At some point, the fixed number of markers on the GigaMUGA will no longer capture increasingly smaller founder haplotype blocks. Missing crossovers significantly reduce QTL resolution, emphasizing the need for alternative approaches are developed and applied.

Low-coverage genotyping-by-sequencing (GBS) followed by SNP imputation is a flexible method for cataloguing segregating genetic variation and performing quantitative genetics. One popular option for GBS is restriction-site associated DNA sequencing (RAD-seq) (Baird et al., 2008), the most popular implementation of which is double-digest RAD-seq (ddRADseq) (Peterson et al., 2012). ddRADseq is commonly used in studies of emerging model systems, where high-quality reference genome resources are absent, and many agricultural species (Elshire et al., 2011), (Sonah et al., 2013), (Furuta et al., 2017), (Fu, 2018), (De Donato et al., 2013), (Wang et al., 2017), (Ivanov et al., 2021). ddRADseq has also been used to study complex trait genetics in outbred rat (Gileta et al., 2020), (Gileta et al., 2022) and mouse populations (Parker et al., 2016), (Gonzales et al., 2018). The main advantages of ddRADseq relative to genotyping arrays are that (1) no prior knowledge of where polymorphisms lie in the genome is required, (2) the reagents required to perform library preparation for this type of sequencing are relatively inexpensive, and (3) genotypes are theoretically captured at the same loci for all individuals in the population, making the translation from genotypes to genetic map less complex.

Another GBS approach is low-coverage whole-genome sequencing (lcWGS). This approach is commonly used in human genetics studies (consortium, 2015), (Homburger et al., 2019), (Li et al., 2021), and is increasingly being adopted in other systems (Ding et al., 2023), (S. Liu et al., 2024), (Lloret-Villas et al., 2023), including mice (Nicod et al., 2016), (Zou et al., 2022). Compared to genotyping arrays, lcWGS offers several experimental advantages. First, variant sites are captured in a relatively unbiased fashion, free of the ascertainment challenges posed by the deliberate selection of target sites profiled on commercial genotyping arrays. Second, imputation algorithms can accurately call millions of variants by inferring latent haplotypes when reference panels are available, offering much higher genetic resolution than arrays. Third, novel genetic variants can be captured, including de novo mutations.

A recent study by Chen et. al. (Chen et al., 2024) demonstrated the performance of both ddRADseq and lcWGS for imputing variants in a large population of outbred rats derived from eight inbred founder strains. While they established the utility and power of both GBS strategies for genome-wide association mapping (GWAS), the question of whether ddRADseq or lcWGS can be used for accurate haplotype reconstruction remains unexplored.

We have implemented an accurate and scalable lcWGS workflow for complex mouse crosses and demonstrate its potential application to the DO population. We performed a comparative evaluation of both ddRADseq and lcWGS and show that both methodologies can be used to reconstruct founder haplotypes of DO animals with high accruacy. Furthermore, we down-sampled alignments produced using each workflow and demonstrated that haplotype reconstructions reflected those produced by the GigaMUGA with high concordance down to 0.1X coverage. We also recovered cryptic haplotype blocks undetected by the GigaMUGA, highlighting the utility of increased genotyping resolution in the future generations of the DO. In a larger independent cohort of embryoid bodies derived from DO embryonic stem cells, we showed that local expression QTLs (eQTLs) are detected with similar power and greater precision with lcWGS than the GigaMUGA and with no clear bias in allele effect size estimation. We conclude that lcWGS can be applied to the DO population and, with simple alterations to our workflow, other complex mouse crosses to accelerate systems genetics by cheaply and accurately obtaining genotypes.

## MATERIALS AND METHODS

### Mouse Samples

We obtained tail tips from 48 J:DO (DO; RRID:IMSR_JAX:009376) mice (24 females and 24 males) (NCBI:txid10090) from outbreeding generation 41 and isolated DNA using an in-house protocol. Briefly, tail samples were lysed in a standard lysis buffer (100 mM NaCl, 10mM EDTA pH 8, 10mM Tris-HCl pH 8, 0.5% SDS, 20 units/ml Proteinase K) at 55°C for 3 hours with intermittent vortexing, followed by incubation for 0.5 hours with RNAse A (0.3 mg/ml). The lysate solution was then incubated with agitation in a 2:4 v/v solution of 2.5 M NaCl and isopropanol for 10 minutes to precipitate DNA. Two volumes of silica-coated superparamagnetic beads (SeraSil-Mag 400, Cytiva) were then added to the precipitated DNA. After 3 minutes incubation to allow DNA binding, the beads were settled on a magnetic tube rack, the supernatant was discarded, and the beads were washed twice with 80% ethanol and allowed to dry briefly. The beads were then resuspended in TE buffer or 10 mM Tris-HCl pH 8 for 3 minutes, and the supernatant containing the gDNA was separated from the beads using the magnetic tube rack. DNA was suspended at a concentration of 1 μl beads/40 μl TE. This DNA was used in both array- and sequencing-based genotyping protocols described below.

### lcWGS Library Protocol

We prepared lcWGS libraries using the plexWell LP384 kit (seqWell, Beverly, MA), according to manufacturer’s instructions. Small gDNA fragments (< 1 kb) were removed from all samples prior to beginning the protocol using AMPure XP Beads (Beckman Coulter, Brea, CA) at 0.4X concentration to reduce sample loss during the post-PCR size selection clean-up. After library preparation, sample pools were bead size-selected to retain fragments > 375 bp in length.

### ddRADSeq Library Protocol

We prepared ddRADseq libraries as previously described (Peterson et al., 2012), (Bayona-Vasquez et al., 2019), with minor modifications. Oligos were designed for *PstI* and *MseI* restriction overhangs, and PCR primers were designed to contain standard i7 indices and a unique molecular identifier (UMI) in place of the i5 index (Hoffberg et al., 2016) (supplementary table with oligo and adapter info). Samples (300 ng each) were pooled with 1 μl of a Pstl and 1 μl of an MseI adapter (both 5 μM stocks) in a 96 well plate so that all samples would receive a unique combination of barcodes. The plate was then covered in AirPore tape (Qiagen, Germantown, MD), and the samples were allowed to air dry at room temperature. DNA was digested using 10 units each of PstI-HF and MseI (New England Biolabs, Ipswich, MA) in 20 μl reactions at 37**°**C for 1 hour. Immediately after RE incubation, 30 μl of prepared T4 DNA Ligase (640 U/reaction; New England Biolabs) in buffer was added to each well. An alternating sequence of ligation-RE digestion alternating sequence was run (22**°**C for 20 minutes, 37**°**C for 10 minutes, repeated twice), followed by exposure to 65**°**C for 20 minutes to heat inactivate the enzymes. Samples were then pooled by column and were cleaned and size-selected to 250-500 bp fragments using AMPure XP Beads. The UMI and i7 adapters were then completed using a double PCR-bead cleanup protocol, as described previously (Hoffberg et al., 2016). We amplified the libraries using 8 PCR cycles.

### Sequencing

Libraries were multiplexed and sequenced on an Illumina NovaSeq 6000 using the S4 Reagent Kit v1.5 to generate 150 bp paired-end reads on an S4 flow cell.

### Microarray-based Genotyping

We sent the DNA to Neogen (Lansing, MI) for genotyping on the GigaMUGA (Morgan et al., 2015). Using reference GigaMUGA data from the DO founder strains, we inferred the haplotype structure of the DO mice using a hidden Markov model in the R/qtl2 software.

### Sequencing data processing

We obtained between 10 and 48 million paired-end sequencing reads for each DO mouse sample. Differences between the lcWGS and ddRAD-seq sequencing library preparation required tailored sequence analysis pipelines.

#### lcWGS

We removed sequencing adapters using fastp (Chen et al., 2018). Reads were aligned to the mouse reference genome build GRCm39 with a mismatch penalty of 0.8 using BWA (version 0.7.17) (https://github.com/lh3/bwa) (https://arxiv.org/abs/1303.3997). We sorted the alignments, removed duplicate alignments using Picard (version 2.26.10) ( https://broadinstitute.github.io/picard/) and determined the sample-specific average genome-wide coverage using the Samtools *depth* command (version 1.16.1)(Li et al., 2009), (Danecek et al., 2021).

#### ddRADseq

We removed sequencing adapters and PCR cloned reads using the *clone_filter* command implemented in Stacks (version 2.64)(Catchen et al., 2011), (Catchen et al., 2013). We aligned and sorted the ddRADseq alignments similarly to the seqWell alignments but did not remove duplicates, as we expect sequences arising from restriction enzyme digests to be roughly duplicated.

### Alignment downsampling

One of the key benchmarks we sought in our study was the lowest coverage with which a sample could be sequenced and still obtain accurate haplotype reconstruction. After obtaining alignments from each preparation method, we used the calculated coverage to down-sample the alignment using the Samtools *view* command with the -s flag, supplying the percentage by which to down-sample the alignment to achieve the desired down-sampled depth.

### Reference data preparation

We prepared reference haplotypes for the DO founders using SNPs from the Mouse Genomes Project VCF on mouse reference genome build GRCm39 (https://ftp.ebi.ac.uk/pub/databases/mousegenomes/REL-2112-v8-SNPs_Indels/). We identified biallelic SNPs segregating among the founders for each chromosome and removed variants which were heterozygous in any founder. The founder genotypes were converted to chromosome-specific IMPUTE2 haplotype, legend, and sample files.

### SNP Imputation

We ran QUILT (version: 1.0.5)(Davies et al., 2021) using the founder reference data and the sample alignment files from each sequencing method. We set the *nGen* parameter to the median generation of all mice in the initial DO population (41) and the shuffle_bin_radius parameter to 2000. We then converted the QUILT-generated VCF file and the filtered founder genotypes for each population to files required by R/qtl2 to perform haplotype reconstruction using the Bioconductor VariantAnnotation package (Obenchain et al., 2014), R/qtl2, and R/qtl2convert (https://github.com/rqtl/qtl2convert).

### SNP grid for haplotype reconstruction

In a set of 48 samples, we demonstrated that haplotypes can be accurately reconstructed, although calculating the founder allele probabilities using R/qtl2 is time intensive. Retaining all imputed variants is also memory intensive. For these reasons, we opted to anchor all imputed SNPs to 1 million SNPs selected with even physical spacing from the Mouse Genome Project VCF (referenced throughout as the “1M variant grid”) prior to haplotype reconstruction.

We used a set of heuristics to filter the SNPs that would enter the founder haplotype reconstructions because SNP imputation quality is directly affected by sequencing coverage, which is a crucial independent variable in our analyses. We retained only SNPs approximately in Hardy-Weinberg equilibrium (P > 0.05), recognizing that many variants in a small population derived from eight inbred founders will likely not meet this assumption. Finally, we retained only SNPs that exceeded a 0.95 information (INFO) score. Often at extremely low coverage levels, few to no SNPs met these criteria. To coerce a comparison between coverage levels, if fewer than 10,000 SNPs on a single chromosome met the 0.95 INFO score cutoff, we reduced the cutoff value by 0.01, collected the SNPs that exceeded the threshold until greater than 10,000 variants could be used, and recorded that INFO score threshold. This produced the final set of filtered sample genotypes used for haplotype reconstruction.

### Haplotype reconstruction

The sample genotypes were used to estimate each of the 36 DO founder genotype probabilities at each filtered imputed marker using a hidden Markov model. Genotype probabilities were condensed to 8-state allele probabilities for each chromosome using the R/qtl2 *genoprob_to_alleleprob* function. Throughout, we refer to these 8-state allele probabilities as haplotype reconstructions. We developed a Nextflow pipeline (Di Tommaso et al., 2017) that executes all steps from sequence data processing to haplotype reconstruction (https://github.com/TheJacksonLaboratory/quilt-nf).

### Comparing haplotype reconstructions

As one benchmark of the performance of the two GBS approaches, we compared the sample allele probabilities obtained from both ddRADseq and lcWGS with those obtained on the GigaMUGA array. We identified the nearest variant in our GBS dataset to each GigaMUGA marker and measured the cosine similarity of the allele probability matrices at that location:

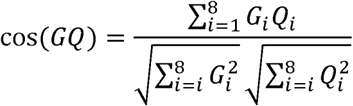

where *G_i_* and *Q_i_* index the allele probabilities for GigaMUGA and GBS genotypes, respectively, for founder *i*. We calculated the mean cosine similarity across all loci as the mean similarity between methods.

#### Formula 1

As a way of prioritizing specific genomic regions in each sample with low concordance, we flagged runs of genotypes with cosine similarity values lower than 0.87. This threshold was selected because it represents the cosine similarity between the following two allele probability vectors, each representing a reconstruction methodology:

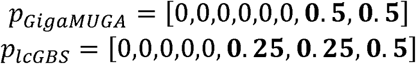

This hypothetical scenario represents a difference in the inferred founder allele probability for one of the two alleles at the locus and signaled to us a sufficient deviation to merit further investigation.

### High-coverage WGS

We selected 10 of the original set of 48 DO samples to be sequenced at high coverage in order to assess the concordance of SNP calls from QUILT. We prepared hcWGS sequencing libraries using the KAPA HyperPrep Kit (Roche Sequencing and Life Science) according to manufacturer instructions. Libraries were sequenced on an Illumina NovaSeq X Plus to generate 150 bp paired-end reads.

### High-coverage WGS data processing

We used the JAX NGS Ops WGS analysis workflow (cs-nf-pipelines; https://zenodo.org/records/15021465) to process our hcWGS sequencing reads. Briefly, adapters were trimmed using fastp and aligned to the mouse reference genome build GRCm39 with a mismatch penalty of 0.8 using BWA (version 0.7.17) (https://github.com/lh3/bwa) (https://arxiv.org/abs/1303.3997). Alignments were sorted using Picard (version 2.26.10) ( https://broadinstitute.github.io/picard/) and SNPs were called using the GATK suite of variant calling tools. SNPs and indels were called using GATK HaplotypeCaller (https://www.biorxiv.org/content/10.1101/201178v3), merged with GATK MergeVcfs, and labeled with filtration thresholds using GATK VariantFiltration. These labels were ultimately used to filter variants that were compared to lcWGS imputed variant set to determine SNP concordance between lcWGS and hcWGS. We retained SNPs that met the following criteria: DP > 10, QD > 2, MQ > 40, FS < 60, ReadPosRankSum > -8, ReadPosRankSum < 8, BaseQRankSum > -8, and BaseQRankSum < 8.

### Crossover counts

We determined whether the number of observed crossovers per mouse was approximately equal to expectation. We performed this analysis using the genotype probabilities obtained from ddRADseq, lcWGS, and GigaMUGA genotypes. We used the *maxmarg* function in R/qtl2 to determine the maximum marginal genotype at each locus in every mouse, and then determined the recombination breakpoints and size of each haplotype using *locate_xo* function in R/qtl2.

### Integrating identity-by-descent and discordant ranges

The DO founder strains include five closely related classical inbred mouse strains that are identical-by-descent (IBD) across many loci (Yang et al., 2011). We examined whether IBD segments (1) could be detected based on the founder reference haplotypes derived from the Mouse Genome Project, and (2) overlapped with regions of founder allele probability discordance in the initial set of 48 DO samples. To identify IBD regions among founders, we applied the *find_ibd_segments* function within R/qtl2 to the lcWGS founder genotypes and lcWGS physical map, specifying a minimum LOD score of 10. We then used the *findOverlapPairs* within IRanges (Lawrence et al., 2013) to determine where IBD regions overlapped among strain pairs. For each sample in our DO cohort, we queried any evidence for founder strain IBD in the discordant range and determined whether the average founder allele probabilities for that sample were consistent with the IBD pattern from the lcWGS data. We then recorded any non-zero founder allele probabilities resulting from both the lcWGS and GigaMUGA genotypes and the original ranges of discordance and IBD. It was often the case that 1) multiple founder pairs demonstrated evidence for IBD at a specific locus, 2) certain founder pairs shared IBD regions more often than others, and 3) discordant regions did not overlap with an IBD locus.

### Embryonic stem cell panel derivation

We modified a previously described protocol for deriving embryonic stem cells by breeding male and female DO mice (aged approximately 6 weeks) and culturing embryonic day 3.5 (E3.5) blastocysts (Czechanski et al., 2014). Samples were initially collected into M2 culture media (Gibco). They were then transferred to plates treated with 0.1% gelatin (Stem Cell Technologies) with feeders and 50% DMEM/F12 (Gibco), 50% Neurobasal medium (Gibco), supplemented with N-2 (Gibco), B27 (Gibco), 1% PenStrep (Gibco), 0.05% BSA (Gibco), 2 mM GlutaMAX-1 (Gibco), 1.5×10-4 M monothioglycerol (Sigma), 1000U/ml LIF (R&D Systems), 1 uM PD0325901 (Stem Cell Technologies), 3 uM CHIR99021 (Stem Cell Technologies), and 2% FBS (Gibco). Cells were weaned off feeders in subsequent passages and once cultures were established (around p6) the 2% FBS was withdrawn. Samples were frozen at approximate densities of 5-7×105 cells/ml at passage numbers in freezing media containing 50% culture media, 40% FBS and 10% dimethyl sulfoxide (DMSO). All ESC samples were transferred to liquid nitrogen holding tanks for long-term storage after controlled freeze (-1°C/min) 24–48 hours at -80°C. All ESC lines were tested for bacterial and fungal growth and for the presence of mycoplasma using a PCR detection system. Cell culture contaminants were not detected.

### Embryonic stem cell genotyping

DNA was collected from each DO ESC line (Qiagen DNeasy Blood and Tissue kit) and genotyped using the GigaMUGA as in *Microarray-based Genotyping* above.

### Embryoid body differentiation

Frozen aliquots of p7-10 DO ESC lines were thawed and grown in 2i/lif for 48 hours. Cells were dissociated with TripLE and counted. Using the Integra Assist Plus robot system, 750 cells/well were planted into 96-well spheroid ultra-low attachment plates (Perkin Elmer) in embryoid body medium (DMEM (Gibco), 15% FBS, 2 mM GlutaMAX-1 (Gibco), 1% PenStrep (Gibco), 1% NEAA, 1% sodium pyruvate, 2-mercaptoethanol and 12ng/mL bFGF). Plates were spun at 1200 rmp for 3 minutes to force aggregation of cells. Cultures were grown for 4 days, changing half the media at day 2.

### RNA sequencing

EBs were pooled for each of 183 DO ESC lines and snap frozen in liquid nitrogen. were stored at -80°C prior to RNA isolation. RNA was extracted using a NucleoMag RNA Kit (Macherey Nagel) and purified with a KingFisher Flex system (ThermoFisher). Library preparation was enriched for polyA containing mRNA using the KAPA mRNA HyperPrep Kit (Rocher Sequencing and Life Science). Paired-end sequencing was performed with a read-length of 150 bp on an Illumina NovaSeq 6000.

### Expression QTL mapping

We obtained DNA from the 183 embryoid bodies derived from DO mice (outbreeding generations 40-45) to demonstrate the feasibility of lcWGS in practice. We performed lcWGS (see *lcWGS Library Protocol* and the lcWGS subsection of *Sequencing data processing*) and GigaMUGA genotyping in parallel. Gene expression counts were obtained following RNA sequencing (above) of the same samples using an established GBRS/EMASE workflow. Briefly, RNA sequencing reads were aligned with Bowtie and counted using EMASE (Raghupathy et al., 2018), thereby quantifying allele-specific expression counts of each gene in an unbiased fashion with respect to the GRCm39 reference genome. We then summed across all allele-specific counts to get total gene expression counts for each gene for all samples. Counts were transformed using the DESeq2’s variance stabilization transformation (Love et al., 2014). Gene expression count distributions across a population can often exhibit distributions that stray far from normality, posing a problem for QTL mapping. To reduce the likelihood of non-normal trait distributions driving biased results for one genotyping method over another, we filtered the variance-stabilized counts to gene counts that exceeded 3000 across all samples and genes with at least 100 unique count values.

After these filtering steps, we retained 17,405 genes for analysis. We then rank-Z transformed the variance-stabilized counts. We performed QTL mapping on each of the rank-Z transformed expression trait using the GigaMUGA and lcWGS haplotype reconstructions generated at four different coverage levels: full coverage, 0.1X, 0.05X, and 0.01X. We used the haplotype reconstructions generated at each coverage level as well as the GigaMUGA to fit a linear model with sex and outbreeding generation as additive covariates. We calculated leave-one-chromosome-out kinship matrices using the R/qtl2 *calc_kinship* function to account for relatedness among individuals. We used the R/qtl2 *scan1* function to fit the following model at each marker:

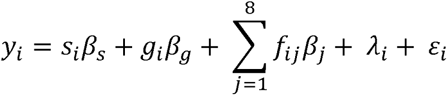

where *Y_i_* is the rank-Z transformed gene expression value for mouse *i*, *β_S_* is the effect of sex, *s_i_* is the sex of mouse *i*, *β_g_* is the effect of outbreeding generation, *g_i_* is the outbreeding generation of mouse *i*, *β_j_* is the effect of founder allele *j*, *f_ij_* is the probability that mouse *i* carries for founder allele *j*, and *λ_i_* is the kinship adjustment for mouse *i.* We identified significant eQTL using the R/qtl2 *find_peaks* function with a LOD threshold of 8, a required peak drop of 5, and a LOD drop of 1.5 to determine the peak width. The distance from each eQTL peak to the midpoint of its respective gene was recorded, and any eQTL located within 2 Mb of the gene midpoint was retained as a local eQTL.

#### Formula 2

We created a 1-to-1 catalog of local eQTLs that were detected in GigaMUGA and lcWGS at each down-sampled coverage level by enumerating the overlapping support intervals for each local eQTL closest to the gene midpoint. For each local eQTL nearest to the gene identified in all 5 eQTL datasets, we quantified the LOD score and the peak width. We also estimated the additive effect of each of the founder alleles for each of these eQTL using the R/qtl2 *fit1* function and calculated the cosine similarity between the GigaMUGA allele effects and each of the down-sampled lcWGS allele effects using Formula 1, but where *G_i_* and *Q_i_*index the allele effect estimates for GigaMUGA and GBS genotypes, respectively, for founder *i* underlying each of the local eQTL.

## RESULTS

### GBS-based haplotype reconstructions are concordant with those from the GigaMUGA

A GBS-based approach for genotyping genetically diverse samples must reproduce haplotypes derived from array-based genotypes to serve as a viable alternative. We benchmarked the performance of two GBS strategies - lcWGS and ddRAD-seq - against GigaMUGA plexWell 384 kit genotypes using a sample of 48 DO mice (see Methods).

For each DO sample, we obtained 18.98 ± 4.77 million mapped high-quality lcWGS reads and 13.97 ± 4.62 million ddRADseq reads (**Figure 1A**), achieving an average genome-wide sequencing depth of 1.14X ± 0.29X and 1.88X ± 0.72X, respectively (**Figure 1B**). The average lcWGS sample aligned to 54.96% ± 8.48% of the genome, while the average ddRADseq samples aligned to only 13.42% of the genome (**Figure 1C**), reflecting the reduced-representation nature of ddRADseq. At sites captured by sequencing reads, we obtained 1.91X ± 0.17X coverage for lcWGS samples and 13.91X ± 3.07X for ddRADseq samples (**Figure 1D**). These sequencing metrics confirmed that we achieved roughly the expected per-site coverage for each library preparation type, i.e. more uniform yet low coverage among lcWGS samples and higher per-site coverage across a smaller fraction of the genome among ddRADseq samples.

**Figure 1.**
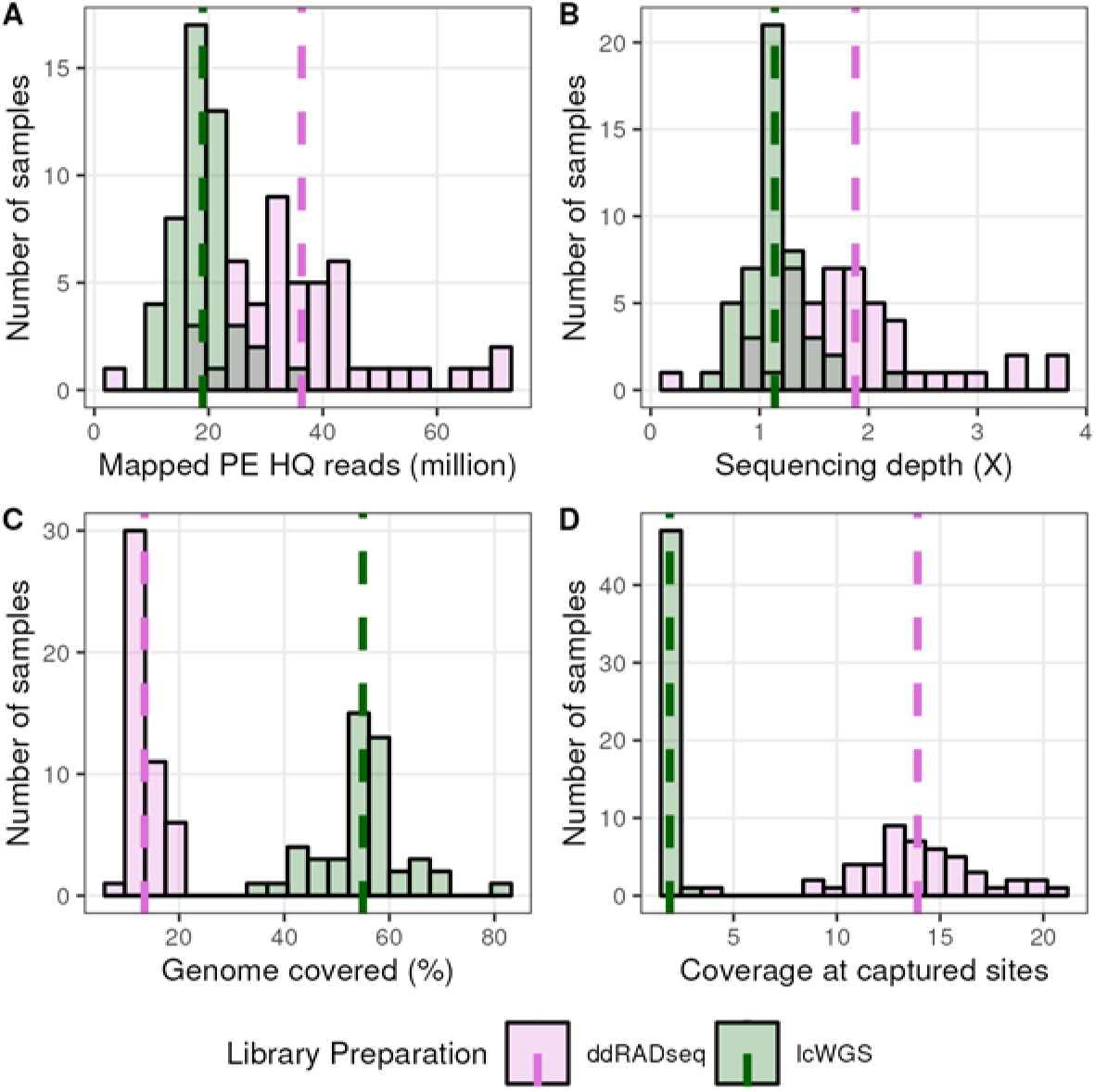
Sequencing statistics of lcWGS and ddRADseq experiments with DO mice. A) Distribution of the number of mapped reads from each sequencing approach for all samples. B) Distribution of the sequencing coverage of all samples. C) Distribution of percent of the genome’s bases covered by sequencing reads among all samples. D) Distribution of the expected coverage by sequencing reads at bases with mapped reads among all samples.

We used our workflow to align the sequencing reads and impute sample genotypes using QUILT (Davies et al., 2021). Each sample was imputed to a resolution of 35,091,075 SNPs originating from reference set of segregating biallelic SNPs in the DO founder strains. We reconstructed haplotypes for each sample based on the founder allele probabilities at each locus using *R/qtl2* using a set of heuristics to filter imputed SNPs (see *Methods*). We used a total of 823,229 filtered SNPs obtained at full coverage from lcWGS samples and 852,154 SNPs from ddRADseq samples (**Figure 2A**). We down-sampled the reads to coverages as low as 0.001X. The number of filtered SNPs used in haplotype reconstruction never dropped below the number of markers on the GigaMUGA. However, at 0.001X coverage, the INFO score threshold for retaining SNPs in haplotype reconstruction was relaxed to average of 0.75 for lcWGS samples and 0.77 for ddRADseq samples (**Figure 2B**) to account for poorer imputation quality.

**Figure 2.**
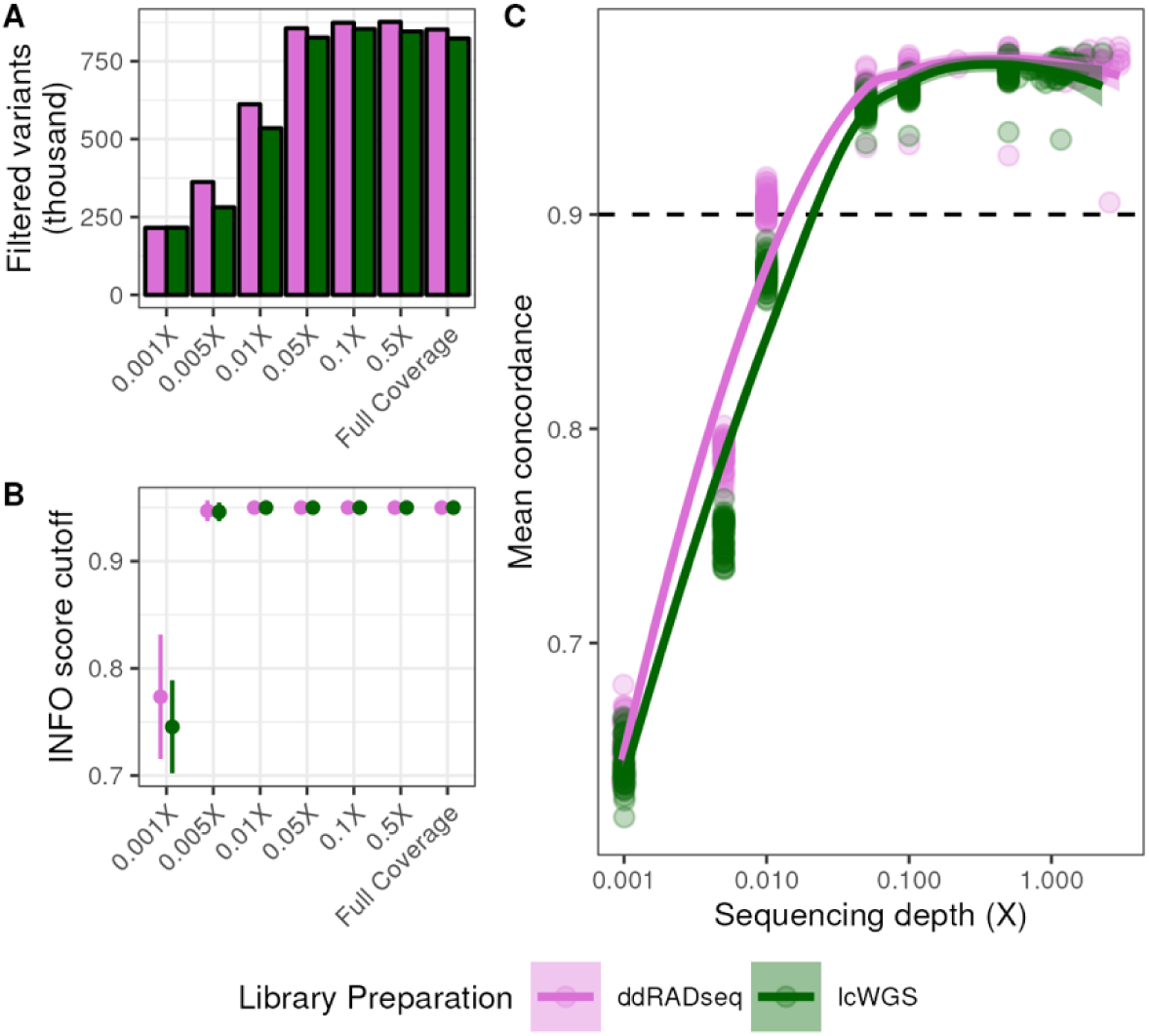
Genotyping-by-sequencing facilitates accurate haplotype reconstruction at low coverage. A) Number of genetic variants retained for haplotype reconstruction after genotype imputation at each coverage level (*see Methods for filtering heuristics*). B) Range of INFO score cutoff values used for filtering imputed SNPs at each coverage level. C) Average sample-level cosine similarity (concordance) of allele probabilities obtained from GBS methods to those obtained from GigaMUGA as a function of sequencing depth.

We then estimated the concordance between the haplotype reconstructions obtained from the GigaMUGA with those obtained from each GBS method. To do this, we interpolated the GigaMUGA founder allele probabilities to the filtered imputed SNP positions, and then calculated the cosine similarity of the eight founder allele probabilities at each position for every sample. We performed this calculation across all down-sampled coverage levels for both lcWGS and ddRADseq to determine the lowest sequencing depth which produced results concordant with the full coverage haplotype reconstructions. The average concordance at full coverage for lcWGS and ddRADseq samples was 0.967 and 0.969 respectively (**Figure 2C**). Concordance rates between lcWGS and ddRADseq did not differ significantly until alignments were down-sampled to 0.1X, at which point the average concordance of lcWGS samples to GigaMUGA was 0.957 while concordance of ddRADseq samples was 0.965 (Two-way ANOVA; Tukey’s HSD *p_adj_* = 0.00008). Down-sampling lcWGS alignments to 0.1X produced a concordance difference from full coverage (Two-way ANOVA; Tukey’s HSD *p_adj_* < 0.000001), resulting in an average concordance of 0.957. Among ddRADseq samples, alignments downsampled to 0.1X did not differ significantly in concordance from full coverage (Two-way ANOVA; Tukey’s HSD *p_adj_* = 0.217). However, those down-sampled to 0.05X were less concordant with GigaMUGA (average concordance value of 0.958; Two-way ANOVA; Tukey’s HSD *p_adj_* < 0.000001). In summary, both lcWGS and ddRADseq produced founder allele probabilities that were highly concordant to those obtained from the GigaMUGA with ultra-low sequencing depth.

We next asked whether either GBS approach estimates the same number of crossovers as the GigaMUGA. We used the *R/qtl2 locate_xo* function to detect crossovers in haplotype reconstructions from both methods and across down-sampling coverages. We found that one sample was a statistical outlier with respect to its crossover count, and for the purpose of comparing methods, it was excluded. At full coverage, autosomal crossover counts obtained using lcWGS and ddRADseq differed significantly from the GigaMUGA (Paired Student’s t-test; *p* < 0.001). Crossover counts obtained from ddRADseq at full coverage were significantly greater than the GigaMUGA by approximately 27.96 autosomal crossovers per mouse, but those obtained from lcWGS were greater by 157.60 autosomal crossovers (**Figure 3A**). Crossover counts obtained from all other coverage levels from lcWGS were also significantly different from the GigaMUGA (Paired Student’s t-test; *p* < 0.05). At coverage levels greater than or equal to 0.05X, lcWGS-based crossover counts exceeded those from GigaMUGA by at least 54 autosomal crossovers per mouse, but lower coverage resulted in underestimation by at least 352 autosomal crossovers per mouse. Meanwhile, ddRADseq-based crossover counts produced with 0.1X coverage were not significantly different from GigaMUGA-based counts (Paired Student’s t-test; *p* = 0.35). Lastly, a theoretical advantage of GBS in the DO is the ability to increase mapping resolution by identifying additional crossovers and smaller haplotype blocks than those inferred with GigaMUGA genotypes. We found that lcWGS (**Figure 3B**) recovered many cryptic haplotype blocks that were not discovered in GigaMUGA and ddRADseq haplotype reconstructions (**Figure 3C**), presumably due to low marker resolution.

**Figure 3.**
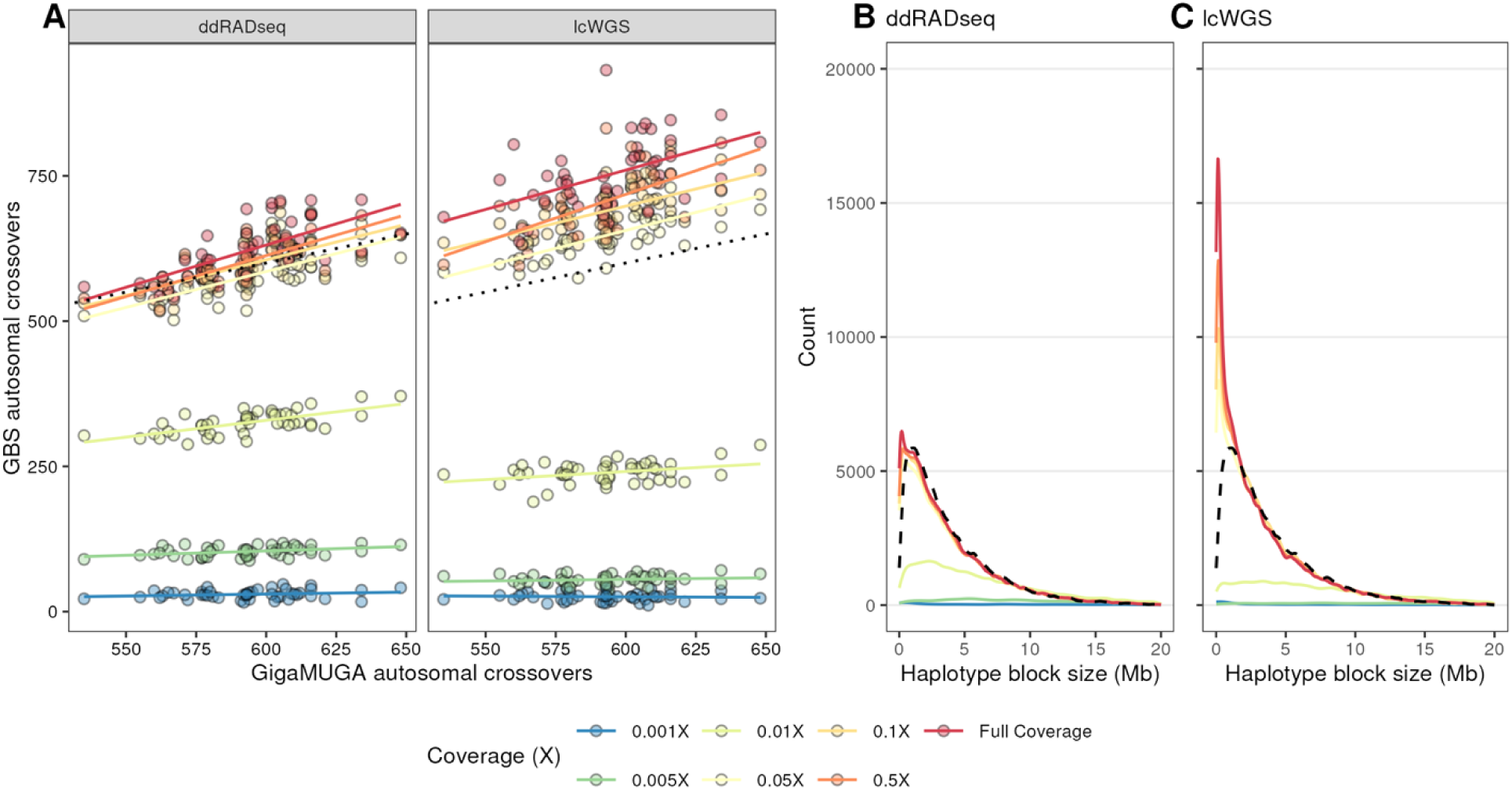
Low-coverage WGS haplotype reconstructions recover additional haplotype blocks in the DO population. A) Autosomal crossover counts from each sample derived from the GigaMUGA (x-axis) compared to each GBS method (facets) across coverage levels. B) Distributions of haplotype block lengths from both GBS methods compared to the GigaMUGA haplotype block size distribution (dashed line).

In summary, we assessed the performance of ddRADseq and lcWGS methodologies as alternatives to the GigaMUGA for genotyping a genetically diverse mouse cohort. Both ddRADseq and lcWGS haplotype reconstructions were concordant with those derived from GigaMUGA genotypes at greater than 0.1X sequencing depth. Both ddRADseq and lcWGS detected more autosomal crossovers compared to the GigaMUGA. However, lcWGS provided a greater increase in resolution than ddRADseq, especially among haplotype blocks less than 1 Mb in length. These results suggest that lcWGS captured the important population-level characteristics of genetically diverse mice more robustly than the ddRADseq protocol we used for these 48 samples.

### lcWGS variant calls are concordant with hcWGS

The existence of well-curated, high-density founder genotypes allows us to impute genotypes with high confidence and hence, to reconstruct the founder haplotypes of DO mice. In practice, investigators may wish to only impute SNPs and proceed with downstream genetic analyses. As an example, the N/NIH heterogeneous stock (HS) rats are an outcross derived from eight inbred strains (Hansen & Spuhler, 1984). To demonstrate the utility of lcWGS for genotyping the HS rat population, the authors sequenced 15,000 rats at low coverage and 88 rats at 33X coverage to compare with their low coverage results and found high concordance (Chen et al., 2024). Studies using the HS rats typically use imputed SNPs for association mapping rather than linkage mapping, so their comparison of lcWGS SNP calls to high coverage WGS (hcWGS) SNP calls is more appropriate in that context. Nevertheless, we sequenced 10 of the DO lcWGS samples at high coverage to determine the concordance of hcWGS variant calls with imputed SNPs derived from lcWGS.

We sequenced the DO hcWGS samples to an 24.28X coverage on average and obtained 30X coverage at 26% of sites. These same samples were sequenced at an average of 1.16X coverage in the lcWGS experiment and their haplotype reconstructions were 97% concordant with the array. We note at this point that, in our haplotype reconstruction workflow, we anchor imputed SNPs derived from lcWGS to a 1M variant grid to expedite haplotype reconstruction in large cohorts. Here, we used the full set of 30,015,813 lcWGS imputed variants that passed the INFO SCORE and HWE filters to compare to hcWGS variant calls. We called 13.46 million ± 0.55 million SNPs per sample with GATK and based on and after filtering hcWGS variant calls (*see Methods*) we reduced the set to 5.24 million ± 0.25 million SNPs. 4.38 million ± 0.22 million of these were common to the lcWGS imputed SNP sites and were compared in genotype status to the original lcWGS samples. Of the common SNP set, 95% were imputed accurately in the same lcWGS samples (**Supplemental Table 1**). These results indicate that not only are lcWGS haplotype reconstructions faithfully replicating those obtained from the GigaMUGA, the imputed variant calls obtained from QUILT match high-quality variants called with hcWGS with high accuracy.

### Identity-by-descent among DO founder strains explains methodological discordance

We observed high genome-wide concordance between array-based and lcWGS haplotype reconstructions with lcWGS coverage as low as 0.1X. We nonetheless sought to characterize areas of the genome with methodological disagreements to understand the limitations of lcWGS for genotyping diverse mouse crosses. Within the filtered variant set for each sample, we flagged regions of the genome where the cosine similarity between allele probabilities among consecutive markers was lower than 0.87 (henceforth referred to as “discordant regions”). At full coverage, this heuristic resulted in an average of 944 discordant haplotypes per individual with an average median length per individual of 0.039 Mb, and average discordance of 0.753 (**Supplemental Figure 1**). Discordant regions were scattered throughout the genome, but with notable hotspots on chromosomes 6, 10, 16, and X (**Supplemental Figure 2**).

Our observation of discordant haplotypes on chromosomes 16 and X aligns with previous reports of depleted genetic variation in similar loci among 12 classical inbred strains (Yang et al., 2009), motivating us to ask the question whether the complex ancestry of inbred mouse strains drives this methodological discordance. To test this hypothesis, we first identified identity-by-descent (IBD) regions between the eight DO founder strains on each chromosome. We then identified discordant regions that overlapped IBD regions by at least 10 kb and pulled the allele probabilities from each genotyping platform through the interval. Using the probabilities that were initially used to calculate concordance, we determined whether founder strains IBD at a given overlapping region were the same founders driving methodological discordance.

77.7% of discordant regions obtained from lcWGS at full coverage overlapped with at least one IBD region. Moreover, of those overlapping regions, 70.8% of discordant regions harbored differing allele probabilities for the founders that were inferred to be IBD (**Figure 4A**). With decreasing coverage, the fraction of discordant regions overlapping an IBD haplotype increased, most apparently at 0.01X coverage and lower. Of the 70.8% of discordant regions putatively driven by IBD among founder strains at full coverage, 93.2% of those haplotypes were IBD among at least two of the classical inbred strains comprising the DO founders (**Figure 4B**). At the chromosome level, the percentage of discordant haplotypes overlapping haplotypes IBD among founder strains identified at full coverage did not differ significantly from the fraction identified at 0.05X coverage (Two-way ANOVA; Tukey’s HSD *p_adj_*= 1) further reflecting overall haplotype reconstruction accuracy at this sequencing depth.

**Figure 4.**
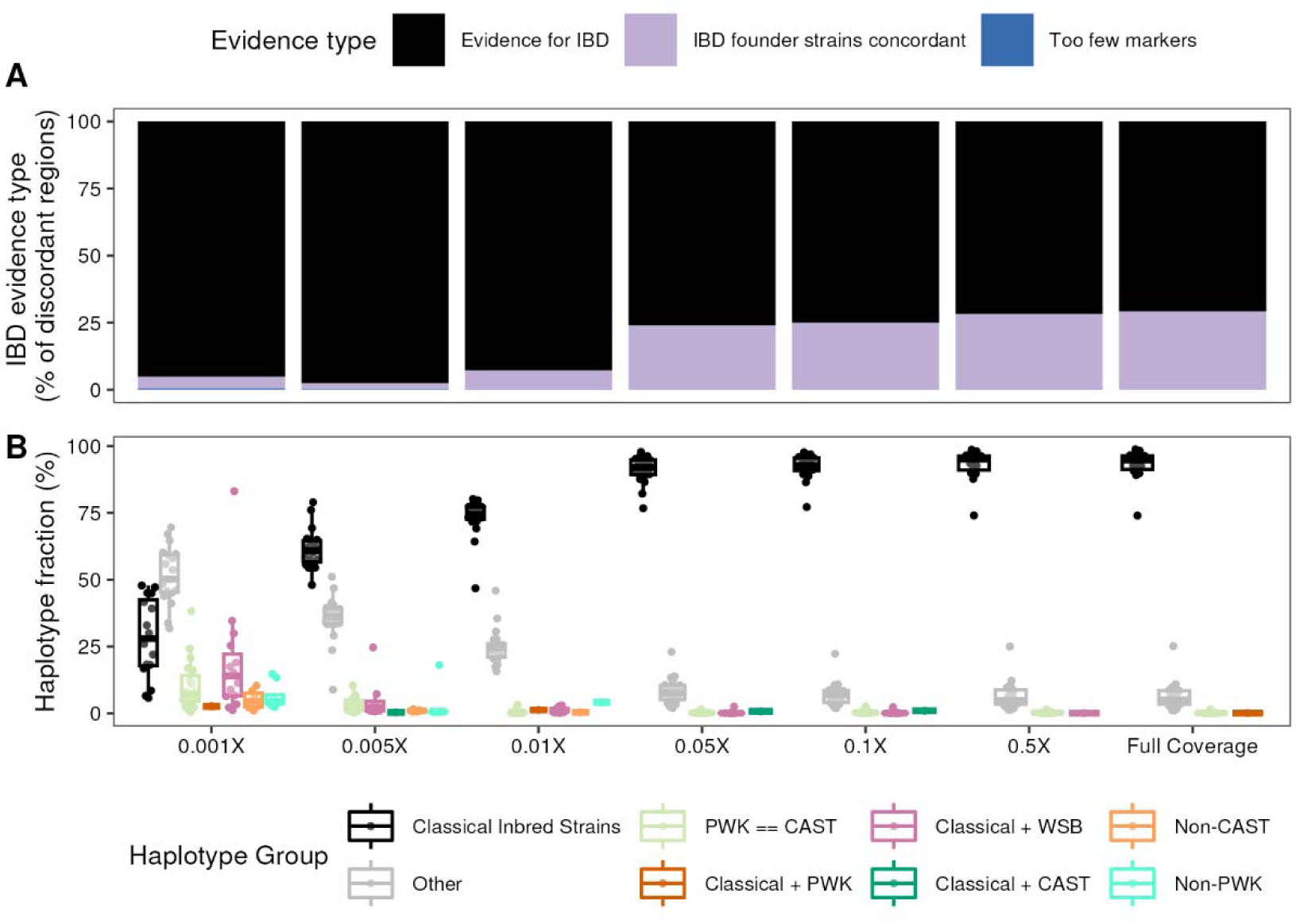
Allele sharing among DO founder strains contributes to methodological discordance. A) Percent of regions with evidence of identity-by-descent (IBD) among any DO founders overlapping discordant regions for which discordant founders are IBD (black) or not (lavender). B) Percent of black regions in A) further categorized into specific allele sharing patterns.

In contrast, at the lowest down-sampled coverage (0.001X), 95.1% of discordant regions harbored differing allele probabilities for the founders that were inferred to be IBD, a full 24.3% higher than full coverage. However, the breakdown of IBD haplotype groups was much more diversified: only 28.2% were IBD among only classical inbred strains, 18.6% were IBD among classical inbred strains and WSB/EiJ, 10.4% were IBD between PWK/PhJ and CAST/EiJ, and 51.8% of IBD haplotypes fell into the category of “Other” which encompasses other inferred haplotype sharing patterns (i.e. sharing between at least one each of a classical founder strain and a wild-derived founder strain). We therefore conclude that loci exhibiting array discordance largely arise from lcWGS allele probabilities modeling real uncertainty in founder haplotype structure due to allele sharing between closely related mouse strains.

### In application: Regulatory variation underlying embryonic stem cell differentiation in the Diversity Outbred population

After benchmarking the accuracy of lcWGS-based haplotype reconstructions, we evaluated the practical performance of lcWGS in a QTL mapping experiment. We differentiated 205 embryonic stem cell (ESCs) lines derived from distinct DO mice into embryoid bodies and collected bulk RNA-seq data for 202 samples. Of these, 183 were also genotyped using GigaMUGA and lcWGS (**Supplemental Figure 3A**). On average, we obtained 17.3 million ± 4.29 million high-quality paired-end reads per sample, resulting in an average sequencing depth of 0.926X ± 0.227X per sample and 2.04X ± 0.156X at captured sites (**Supplemental Figure 3B**). We then used both lcWGS and the GigaMUGA to genotype every sample and performed haplotype reconstruction as outlined above for both genotyping platforms. We used EMASE (Raghupathy et al., 2018) to quantify the transcriptome of each sample, and after quality filtering transcripts (*see Methods*), used a total of 17,405 gene counts as quantitative traits in eQTL mapping.

We mapped a total of 5,509 eQTL using GigaMUGA and from 4,464 to 7,089 eQTL using lcWGS with increasing sequencing depth. Local eQTLs are often robust and reproducible across cohorts of the same population, making them desirable for benchmarking the performance of different genotyping methodologies. For each gene, we selected the closest eQTL to facilitate one-to-one comparisons of local eQTLs between genotyping approaches. Using this heuristic, we reduced the number of local eQTLs identified using GigaMUGA to 4,974. Depending on sequencing depth, between 79% (0.01X) and 99% (0.1X) of these eQTL were also detected using lcWGS. 3,375 GigaMUGA eQTL (68%) were detected across all coverage levels (**Figure 5A**). We used this subset of consistently detected local eQTLs to characterize the effect of sequencing depth on eQTL position, resolution, and effect size between lcWGS and GigaMUGA.

**Figure 5.**
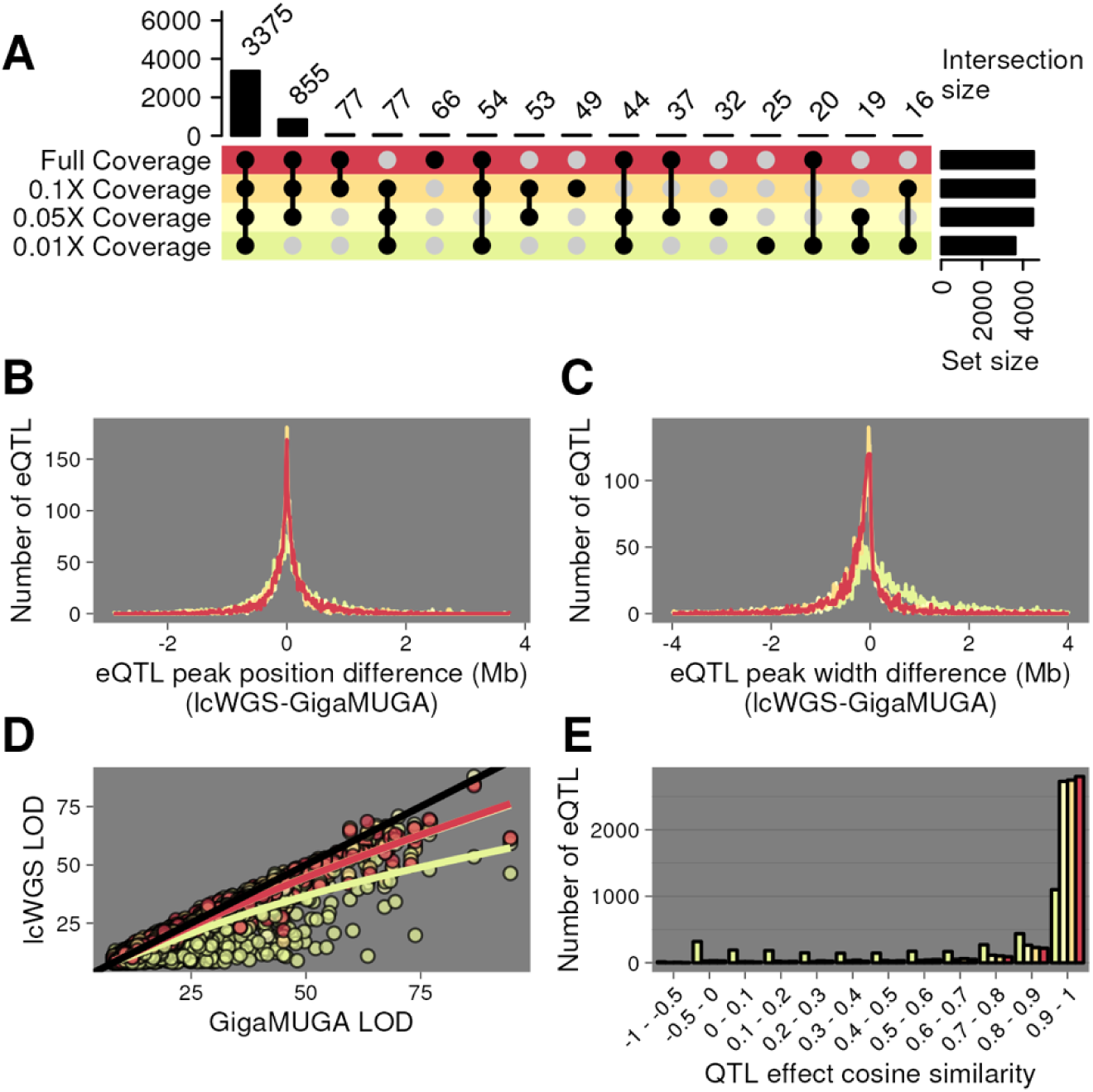
Expression QTL mapping with lcWGS matches or exceeds performance of GigaMUGA. A) UpSet plot (Gu et al., 2016) of local eQTL mapped in the DO (*n* = 183) using GigaMUGA haplotype reconstructions that were also recovered by lcWGS at full coverage (red), 0.1X (pale orange), 0.05X (pale yellow), or 0.01X (pale green). B) Distribution of the difference in peak positions of local eQTL recovered in all lcWGS coverage levels and GigaMUGA. C) Distribution of the difference in 1.5 LOD-drop support interval width of local eQTL recovered in all lcWGS coverage levels and GigaMUGA. D) Scatter plot of the LOD scores of local eQTL recovered with the GigaMUGA (x-axis) and all lcWGS coverage levels (y-axis). E) Histogram of the cosine similarity of additive allele effects underlying local eQTL recovered using the GigaMUGA and all lcWGS coverage levels.

The peak position for each compared eQTL was nearly identical (**Figure 5B**). On average, full-coverage lcWGS eQTL peaks (1.5 LOD-drop support interval) were 0.17 Mb narrower than their respective GigaMUGA eQTL peaks, a difference significantly different from 0 (Student’s t-test, *p* = 0.0007) (**Figure 5C**). We observe no significant difference in peak widths with down-sampling to 0.1X or 0.05X (One-way ANOVA; Tukey’s HSD *p_adj_* ≥ 0.935). At full coverage, lcWGS and the GigaMUGA eQTL LOD scores had a correlation of 0.98. The magnitude of this correlation decreased to 0.88 when lcWGS was downsampled to 0.01X coverage (**Figure 5D**). Finally, we asked whether lcWGS- or GigaMUGA-based haplotype reconstructions yield similar effect size estimates to each founder allele. We extracted the eQTL additive allele effects from each of the 3,375 shared eQTL and calculated the cosine similarity of the coefficients between each coverage level of lcWGS and GigaMUGA. At greater than or equal to 0.05X coverage, at least 82% percent of the shared eQTL had allele effect cosine similarity values greater than 0.9 (**Figure 5E**), demonstrating that there was limited bias in allele effect estimation across detected eQTLs. We conclude that standard QTL analysis in DO mice using GigaMUGA haplotype reconstructions was recapitulated by lcWGS at coverage levels > 0.1X.

## DISCUSSION

As large-scale phenotyping and multi-omics studies become the standard for rigorous systems genetics, the cost and scale of genotyping technologies must adapt to meet the dual needs for cost efficiency and throughput. The DO mouse population is a powerful model for systems genetics, featuring segregating genetic variation from biomedically relevant mouse strains and fine recombination block structure. This structure is largely captured by the GigaMUGA, which remains a robust and reliable genotyping array for mouse crosses. The MUGA-series platform has been the most popular choice for genotyping complex mouse crosses because they have the advantage of assaying the same markers in every cross, which allows for a standardized analysis pipeline. However, as sequencing costs have decreased, there are now less expensive genotyping options which offer increased marker resolution. This is particularly important as multi-generation outcrosses acquire more crossovers and develop finer haplotype block structure. Here, we have shown that lcWGS recovers small haplotype blocks that elude detection by lower resolution genotyping approaches. We found that DO haplotypes observed in the GigaMUGA were consistently recovered by lcWGS reconstructions down to a coverage of 0.1 X, consistent with findings of other groups (Chen et al., 2024).

In this study, we reconstructed DO haplotypes both to compare them to the GigaMUGA-derived haplotypes and to perform eQTL mapping in a DO-derived *in vitro* system. We used concordance with GigaMUGA-derived haplotypes as a reference to evaluate the performance of our SNP imputation. Our approach also enabled us to validate our method using empirical expression data and to test whether founder allele effects were estimated consistently, independent of genotyping platform. Both sample-level concordance and locus-level allele effect concordance were high, suggesting that our lcWGS genotyping approach is capturing the haplotype structure of DO mice. Since linkage mapping using GigaMUGA-derived haplotypes is the most common approach to QTL mapping in the DO, we opted to compare the lcWGS methods to the GigaMUGA rather than hcWGS.

We also compared the number of crossovers per mouse between genotyping methods. In the DO, we expect an average increase of 23.9 observed crossovers per mouse at each outbreeding generation (Broman, 2012), (Gatti et al., 2014). This estimate accounts for crossovers within homozygous regions in DO mice which we cannot observe. Previously, we found that DO mice at generation 11 had approximately 400 crossovers (Gatti et al., 2014). Extrapolating to generation 41 (the outbreeding generation of the 48 DO mice in this study), we would expect approximately 1,117 crossovers, assuming no crossover interference or erosion of *Prdm9* binding sites (Baker et al., 2015). We observed 760 ± 85.8 crossovers per mouse using lcWGS, which is short of the theoretical expectation. Nonetheless, we do recover more short haplotype blocks using lcWGS than with the GigaMUGA. However, we may be missing haplotype blocks due to regions of sequence identity between the DO founders, which lowers the confidence of the haplotype reconstruction model and may render crossovers invisible in genetic data.

We compared ddRADSeq to lcWGS and selected lcWGS based on three main criteria: usability, flexibility, and cost. With regard to usability, the lcWGS reagents are sold as a kit, whereas the ddRADSeq reagents must be purchased separately, streamlining purchasing and laboratory inventory tracking. The lcWGS protocol also requires less hands-on time and fewer steps than the ddRADSeq protocol. On flexibility, the lcWGS protocol is already able to adapt to differences in haplotype block size of DO cohorts over time. ddRADseq is theoretically well-suited for identifying tag SNPs for each haplotype block (Gileta et al., 2020), but we found that ddRADseq missed crossovers and generated fewer haplotype blocks less than 1 Mb than lcWGS. The practical haplotype block detection limit is probably due to a few factors. First, the landscape of RE digest sites in the cohort is specific to the restriction enzymes selected and the size selection range of the restriction fragments. If these parameters are tuned correctly, it may be possible to recover more resolution, but the steps outlined would need to be optimized, lowering reproducibility of genotyping results across experiments. Using lcWGS, one can simply sequence to higher coverage to obtain better resolution as the DO accumulates more crossovers. With regard to cost, all-in price points for ddRADSeq and lcWGS are comparable.

Taken together, we recommend that lcWGS be adopted as the standard for genotyping complex mouse crosses. In order to facilitate the transition from GigaMUGA-based to low-pass GBS, we have assembled our alignment and haplotype reconstruction analysis pipeline into a publicly-available, reproducible Nextflow workflow (Di Tommaso et al., 2017).

Looking forward, we envision several applications and improvements to our approach.

### Modifier screens

A powerful experimental design for detecting genetic modifiers is to identify a strain background or specific mutation that displays the phenotype, and cross that strain to many recombined individuals (Hackett et al., 2022), (Kim et al., 2023), (Gurdon et al., 2024). This crossing scheme offers high resolution mapping when combined with DO mice. Genotyping this number of individuals on the GigaMUGA could be cost-prohibitive, our approach could lower the cost of this experiment drastically because we achieve high haplotype reconstruction accuracy at 0.1X coverage.

### Genetic quality control

The MiniMUGA (Sigmon et al., 2020) is an array with high utility because probes detect commonly engineered mutations and distinguish closely related mouse substrains. Rather than using the MiniMUGA, investigators could use lcWGS to reconstruct haplotypes to verify that the primary and donor strain backgrounds can be distinguished during the generation of mouse mutants, chromosome substitution strains, or introgression lines. Our pipeline is easily tunable for different mouse strains and cross types because reference haplotypes are derived from public data, and most file types are standardized.

### Wild mouse GWAS

The CC and DO carry genetic contributions from three wild-derived strains representing the three major mouse subspecies. However, these backgrounds represent only a fraction of the segregating genetic diversity in wild mouse populations. Opportunities to conduct quantitative genetics will only grow as efforts to catalog wild mouse genetic variation continue. Standard variant calling workflows are sufficient to fuel a standard GWAS in wild mice provided there is ample power. But once high-quality reference haplotypes exist for a subset of wild mice representing major populations, our workflow is well-suited to detect ancestral haplotype structure and link variation in important traits back to these populations using admixture mapping.

### Pan-genome and T2T alignment

In this project, we aligned reads from DO mice to the mouse reference genome, which is based on the C57BL/6J genome. However, the DO founder strains contain considerable genetic variation which is not found in the reference genome (Keane et al., 2011), including large scale structural variants (Yalcin et al., 2011), (Ferraj et al., 2023). Ultimately, we anticipate the need to align reads to a mouse pan-genome (Frankish et al., 2023) based on telomere-to-telomere (T2T) assemblies (J. Liu et al., 2024), (Francis B, 2024).

In conclusion, we demonstrated that lcWGS can be used to genotype DO mice and reconstruct accurate founder haplotypes in a QTL mapping framework. These genotypes were obtained at a fraction of the price of array genotypes and produced not only equivalent mapping results, but on occasion, narrower QTL intervals when benchmarked with empirical data. In the future, we hope to extend our framework to other mouse populations and other types of sequencing libraries, especially given the recent increases in accuracy of long-read sequencing. We hope that our methodology provides a roadmap by which experiments using complex mouse crosses become more accessible and scalable.

## ACKNOWLEDGEMENTS

We gratefully acknowledge the contribution of the Genome Technology Service at The Jackson Laboratory for expert assistance with the work described in this publication. We also gratefully acknowledge the contributions of the Information Technology platform at The Jackson Laboratory (Farmington, CT) for providing and supporting work on the HPC cluster. We would also like to kindly thank Sabina Gude and seqWell, Inc for correspondence and expertise in library preparation optimization.

## DATA AVAILABILITY

Whole-genome sequencing data has been deposited in the NCBI Sequence Read Archive under project numbers PRJNA1259049 (48 DO lcWGS samples), PRJNA1259492 (48 DO ddRADseq samples), and PRJNA1261999 (183 DO lcWGS samples for eQTL mapping). The workflow developed to process sequencing reads, impute genotypes, and perform haplotype reconstruction can be found at https://github.com/TheJacksonLaboratory/quilt-nf. Processed data is available for download at DOI: 10.6084/m9.figshare.28661030.

## STATEMENTS AND DECLARATIONS

The authors declare no competing interests.

## COMPLIANCE WITH ETHICAL STANDARDS

All mouse procedures were reviewed and approved by The Jackson Laboratory’s Institutional Animal Care and Use Committee (protocol #20030).

## SUPPLEMENTAL MATERIALS

**Supplemental Table 1.**
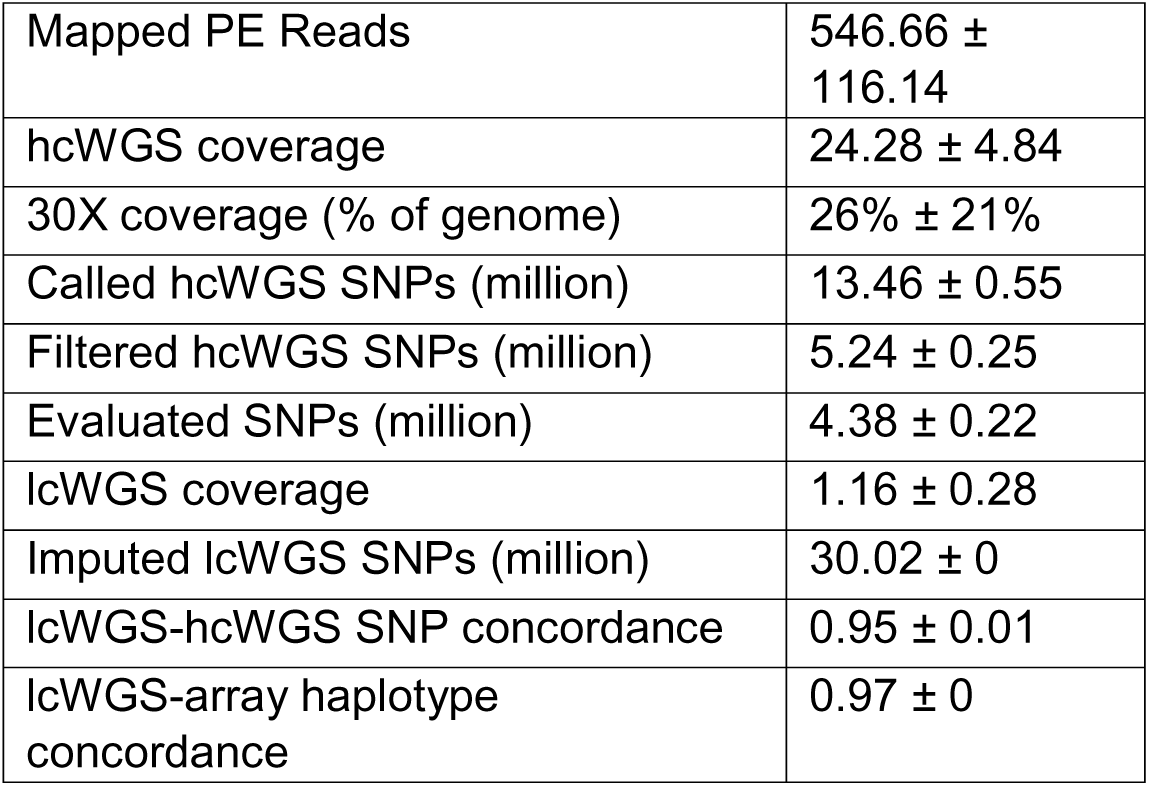
Summary metrics of hcWGS experiment (*n* = 10), SNP concordance, and haplotype concordance of lcWGS in the subset of samples subjected to hcWGS.

**Supplemental Figure 1.**
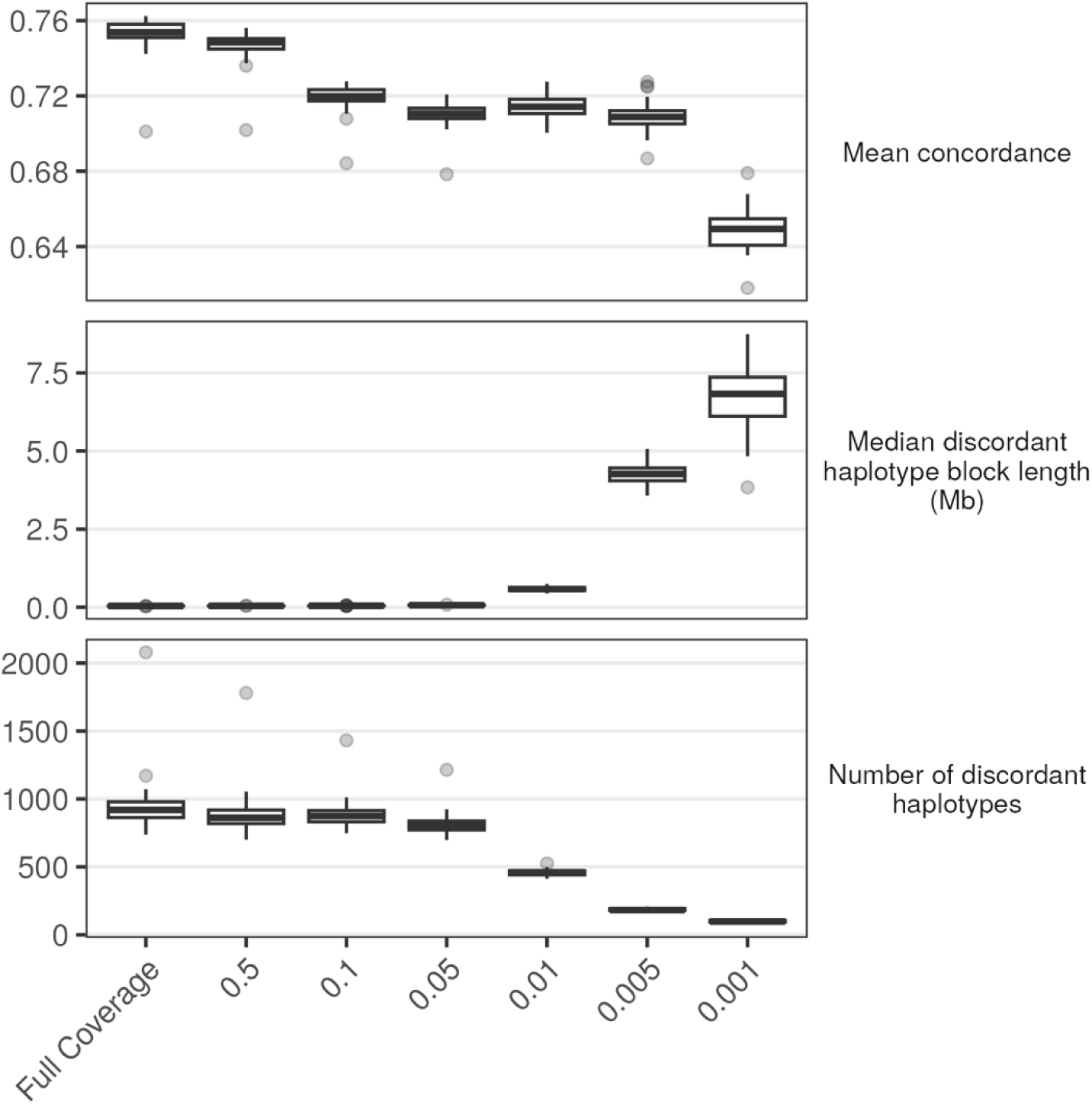
Characteristics of lcWGS haplotype blocks with concordance lower than 0.87 when compared to GigaMUGA haplotype blocks in the same sample at varying coverage specifications.

**Supplemental Figure 2.**
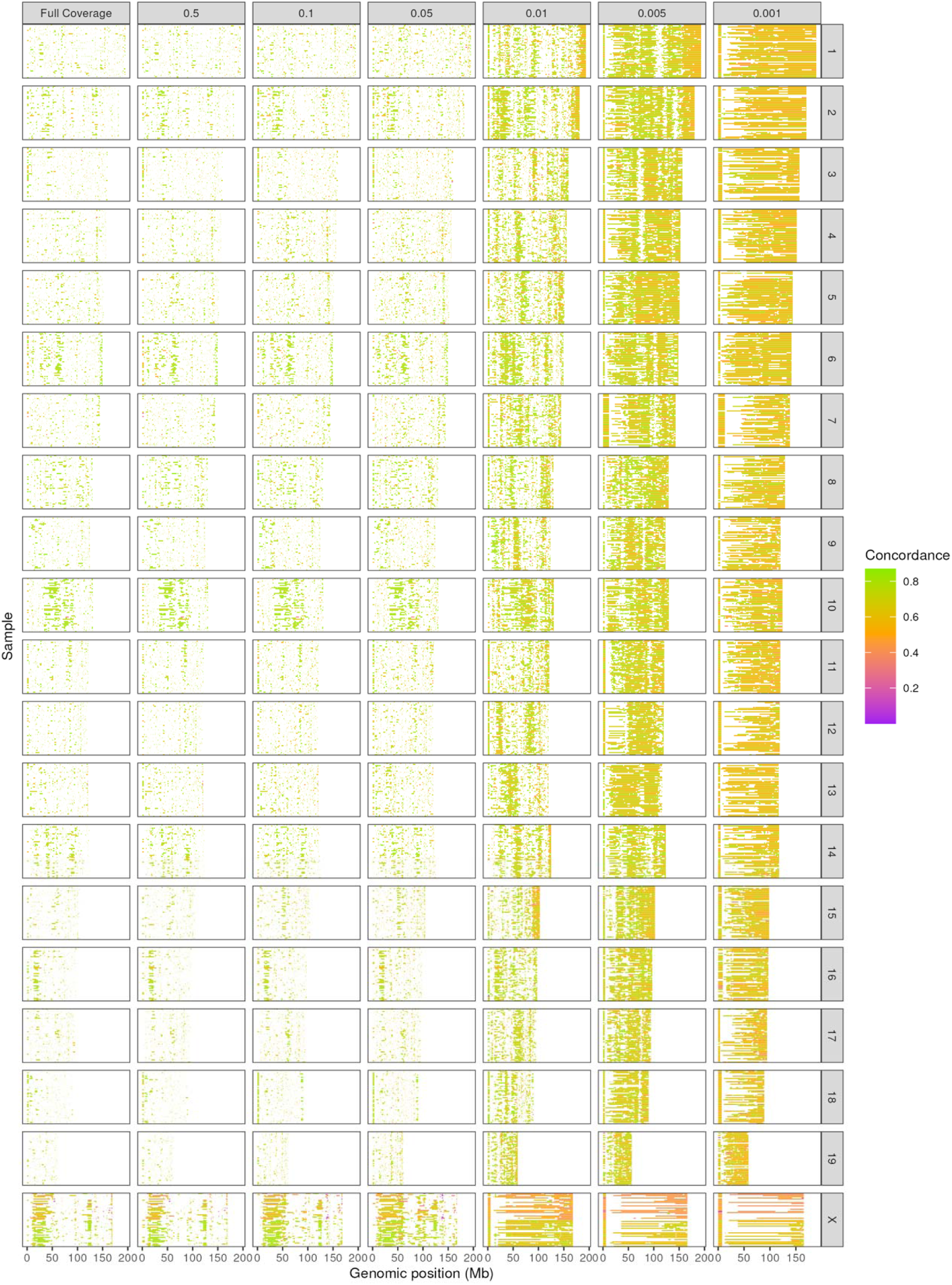
Map of allele probability concordance between lcWGS and GigaMUGA haplotype reconstructions. Each horizontal stripe within each facet represents a region of concordance lower than 0.87 for a given sample, chromosome (vertical facets), and sequencing coverage specification (horizontal facets) combination.

**Supplemental Figure 3.**
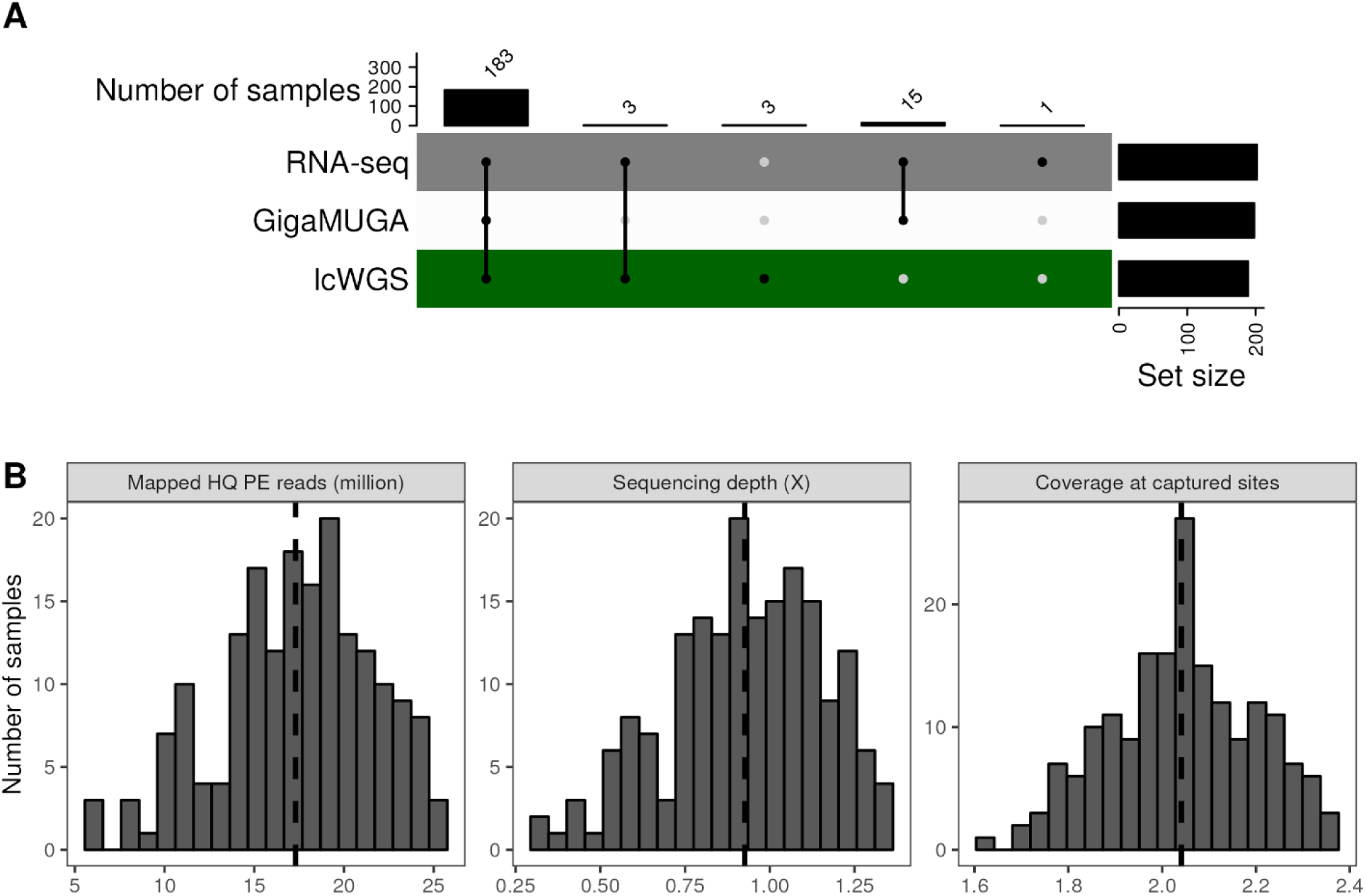
Overview of sample composition and sequencing metrics of DO embryoid bodies (EBs) used for expression QTL analysis. A) UpSet plot (Gu et al., 2016) of DO EBs and data types obtained. B) Sequencing metrics for 183 DO EBs used for lcWGS.

